# Gene expression analysis provides insight into the physiology of the important staple food crop cassava

**DOI:** 10.1101/073213

**Authors:** Mark C. Wilson, Andrew M. Mutka, Aaron W. Hummel, Jeffrey Berry, Raj Deepika Chauhan, Anupama Vijayaraghavan, Nigel J. Taylor, Daniel F. Voytas, Daniel H. Chitwood, Rebecca S. Bart

## Abstract

- Cassava (*Manihot esculenta*) feeds approximately 800 million people worldwide. Although this crop displays high productivity under drought and poor soil conditions, it is susceptible to disease, postharvest deterioration and the roots contain low nutritional content.
- Here, we provide molecular identities for eleven cassava tissue types through RNA-sequencing and develop an open access, web-based interface for further interrogation of the data.
- Through this dataset, we report novel insight into the physiology of cassava and identify promoters able to drive specified tissue expression profiles. Specifically, we focus on identification of the transcriptional signatures that define the massive, underground storage roots used as a food source and the favored target tissue for transgene integration and genome editing, friable embryogenic callus (FEC).
- The information gained from this study is of value for both conventional and biotechnological improvement programs.

## Introduction

Cassava (*Manihot esculenta*) is the food security crop that feeds approximately 800 million people worldwide (Liu *et al.*, 2011; Howeler *et al.*, 2013). Although this crop displays high productivity under drought and poor soil conditions, it is susceptible to disease, postharvest deterioration and the roots contain low nutritional content (Gegios *et al.*, 2010; Stephenson *et al.*, 2010; Patil *et al.*, 2015). Cassava improvement programs are focused on addressing these constraints but are hindered by the crop’s high heterozygosity, difficulty in synchronizing flowering, low seed production and a poor understanding of the physiology of this plant (Ceballos *et al.*, 2004). Among the major food crops, cassava is unique in its ability to develop 51 massive, underground storage roots. Despite the importance of these structures, their basic physiology remains largely unknown, especially the molecular genetic basis of storage root development. Similarly, in cassava, the favored target tissue for transgene integration and genome editing is a friable embryogenic callus (FEC) (Taylor *et al.*, 2001; Bull *et al.*, 2009; Taylor *et al.*, 2012; Zainuddin *et al.*, 2012; Nyaboga *et al.*, 2013). Little is known concerning gene expression in this tissue, or its relatedness to the somatic organized embryogenic structures (OES) from which it originates (Gresshoff & Doy, 1974; Taylor *et al.*, 2012; Chauhan *et al.*, 2015). Here, we provide molecular identities for eleven cassava tissue types through RNA-sequencing and develop an open access, web-based interface for further interrogation of the data. Through this dataset, we report novel insight into the physiology of cassava and identify promoters able to drive specified tissue expression profiles.

## Materials and Methods

### Plant material and tissues sampled

Plant tissues were sampled from 3-month-old TME 204 cassava plants, grown in a greenhouse at the Donald Danforth Plant Science Center (St. Louis, MO). Plants were established from in vitro micropropagated plants (Taylor *et al.*, 2012) and grown in a 12 h light:12 h dark photoperiod (250-500 µmol s^-1^ m^-2^ irradiance Daytime day, temperatures ranged 69 from 28°-32°C with 70% relative humidity, and night time, temperatures ranged from 25°-27°C with 70% relative humidity. The following tissues were sampled from these plants at 2pm: leaf blade, leaf midvein, petiole, stem, lateral buds, shoot apical meristem (SAM), storage roots, fibrous roots, and root apical meristem (RAM). For non-meristem tissues, approximating 100 mg of tissue was collected in 3 separate biological replicates. For the SAM and RAM, 6 meristems were dissected and pooled for each of 3 biological replicates. All samples were frozen in liquid nitrogen after collection. Samples of TME 204 OES and FEC were generated as described previously (Chauhan *et al.*, 2015). The OES induced on DKW/Juglans basal salts (*Phyto*Technology Laboratories, Kansas, USA) containing Murashige and Skoog (MS) vitamins and supplemented with 2% w/v sucrose, 50 µM picloram was sampled after four weeks of culture. The OES was separated from the non-embryogenic tissues and collected in 2ml sampling tubes. FEC tissues were sampled after three weeks of culture on Gresshoff and Doy basal medium supplemented with 2% w/v sucrose, 50 µM picloram, 500 µM tyrosine and 50 mg/l moxalactam. Approximately 200 to 250 mg of tissue was collected and the tubes containing the tissues were immediately placed on dry ice.

### Preparation of RNA-seq libraries and Illumina sequencing

For non-meristem tissues, total RNA was isolated with the Spectrum Plant Total RNA Kit (Sigma). For SAM and RAM tissues, total RNA was isolated with the Arcturus PicoPure RNA Isolation Kit. RNA quality was assessed on an Agilent Bioanalyzer. For library preparation with tissues other than SAM and RAM, 5 µg of RNA was used as input. For SAM and RAM, six samples were pooled to obtain a total of 500-600 ng each. The NEBNext Poly(A) mRNA Magnetic Isolation Module (New England BioLabs) was used to isolate mRNA, which was then used for library prep using NEBNext mRNA Library Prep Master Mix Set for Illumina (New England BioLabs) with 13 cycles of PCR amplification. Standard library prep protocol was followed for all samples, except for the SAM and RAM in which 1 µL of fragmentation enzyme was used instead of 2 µL, and 0.5 µL of random primer instead of 1 µL. Library quality was assessed with the Agilent Bioanalyzer. In total, 32 RNA-seq libraries were made from 11 different tissue types with 3 biological replicates each, except for storage root which only had 2 biological replicates. All libraries were multiplexed into one lane of Illumina HiSeq2500.

### Read mapping and gene expression analysis

Illumina RNA-seq reads from each replicate were cleaned using Trimmomatic version 0.32 (Bolger *et al.*, 2014). Using TopHat2 version 2.1.0 (Trapnell *et al.*, 2009), these cleaned reads were then mapped to the version 6.1 draft assembly of *Manihot esculenta* AM560-2 provided on Phytozome10.3 (http://phytozome.jgi.doe.gov/pz/portal.html). The read mapping output was linked to candidate gene models for each sample using Cufflinks version 2.2.1 (Trapnell *et al.*, 2010). The gene models from all samples of the experiment were merged into one gene model file using Cuffmerge version 2.2.1. Using the output from Cuffmerge and the read mapping files from each replicate, a differential expression analysis between tissue types was performed using Cuffdiff version 2.2.1. Quality checks were performed on the Cuffdiff output using cummeRbund version 2.6.1 in R (R Core Team, 2015). The output of Cuffdiff was processed in Python with the pandas, numpy, and seaborn packages to visualize the expression data (McKinney, 2010).

### Multivariate statistics

Analysis of transcript expression profiles began with those transcripts with 1) FPKM values above a threshold of 1 FPKM and 2) those transcripts significantly differentially expressed in at least one pairwise comparison of all tissues types against each other. For Principal Component Analysis (PCA) of tissue replicates (**Fig. 3A-B**), the prcomp() function in R was used with scaled FPKM values across transcripts as input. For the PCA of transcript profiles (**Fig. 3C-D**), the prcomp() function was again used with scaled mean FPKM values across tissues as input. Self-Organizing Maps (SOMs) were performed using the Kohonen package in R. Scaled mean transcript expression values across tissues were assigned to four nodes in a 2x2 hexagonal topology over 100 training iterations. To focus on those transcripts with expression profiles closest to over-represented patterns of variance in the dataset, only those transcripts with distances from their respective nodes less than the median for the overall dataset were subsequently used and projected back onto the transcript PCA space. Data visualization for the above was carried out with the ggplot2 package in R, using geom_point() and geom_line() functions, among others. The color space for the above was determined using palettes from colorbrewer2.org.

### Gene Ontology (GO) enrichment analysis

GO enrichment analysis was completed using the Python goatools package (https://github.com/tanghaibao/goatools). goatools was run with the --fdr flag to calculate the False Discovery Rate (FDR) error corrected p-value, and the --no_propagate_counts flag to prevent nodes at the root of the GO tree from being included in the analysis. GO terms for each gene were used from the annotation file provided on Phytozome. GO enrichment output was then filtered to include only enriched GO terms with a FDR error corrected p-value < 0.001. For SOM node GO enrichment, each SOM node identified above was processed separately. The genes identified as part of the SOM node were used as the study group, and all genes expressed greater than 1 FPKM in at least one tissue with significant differential expression in at least one pairwise comparison were used as the population or background group. For pairwise tissue comparison GO enrichment, genes identified as significantly upregulated with a |log_2_(fold_change)| > 2 in one tissue were treated as one study group to look at each tissue separately. This resulted in two GO enrichment analyses for each pairwise comparison. Genes with at least 1 FPKM in either tissue were used as the background dataset.

### Identification of genes with strong, constitutive, and tissue-specific expression patterns

Custom Python code was used in a Jupyter notebook using the Pandas, NumPy, Seaborn, and SciPy packages to organize, process, and display the data **(SuppFile3)**. Genes with strong expression across all tissue types were identified using expression values from the gene_exp.diff file produced by Cuffdiff. The genes were first checked for functional annotations, then shortened to a list of genes with a minimum expression of 300 FPKM in each tissue sampled. Specifically and constitutively expressed genes were identified using expression values from each replicate in the genes.read_group_tracking file produced by Cuffdiff. Genes used were annotated in the AM560-2 v6.1 assembly on Phytozome10.3. For specifically expressed genes, this list was then subset by selecting genes with expression greater than 10 FPKM in the tissue(s) specifically expressing the gene, and no more than 1 FPKM in all other tissues. For a more relaxed analysis, genes were required to be expressed greater than 8 FPKM in the tissue(s) specifically expressing the gene, and no more than 4 FPKM in all other tissues. Constitutively expressed genes were identified using the replicate expression data. This list was filtered to include only genes with greater than 40 FPKM in all replicates, and then the coefficient of variation was calculated across all replicates for each gene.

### Data availability

A graphical user interface was created using R Shiny (v 0.13.2) to explore the tissue-specific data and discover trends therein. This application uses data from RNA-seq differential expression analysis completed with the Tuxedo Suite pipeline (v 2.2.1), functional gene annotations from the Joint Genome Institute’s Phytozome, and analysis from principle components (prcomp in R “stats” package v 3.2.3) and self-organizing maps (som in R “kohonen” package v 2.0.19). The application has two main features: 1) gene discovery based on gene expression patterns across tissues, and 2) creation of a tissue-specific heatmap of known or newly discovered genes for visualizing expression patterns. Detailed instructions are included in the application. The application can be found at: shiny.danforthcenter.org/cassava_atlas/. Additional R packages used in this application include: png (v 0.1-7), grid (v 3.2.3), ggplot2 (v 2.1.0), shinyBS (v 0.61), shinydashboard (v 0.5.1), DT (v 0.1), stringr (v 1.0.0), mailR (v 0.4.1), and shinyjs (v 0.5.2).

### In planta expression assays

Promoter fragments, listed in SuppFile4, were cloned from cassava variety TME419 into a pCAMBIA vector upstream of GUS. Constructs were transformed into *Agrobacterium tumefaciens* strain LBA4404. Strains carrying the reporter constructs were re-suspended in IM media (10 mM MES, pH5.6; 10 mM MgCl_2_; 150 µM Acetosyringeone), incubated at room temperature for 3 hours and then infiltrated into *Nicotiana benthemiana* leaves at an OD600 = 0.1. 48 hours post inoculation, leaves were detached and placed in a petri dish. GUS staining solution (0.1 M NaPO_4_ pH7; 10 mM EDTA; 0.1% Triton X-100; 1 mM K_3_Fe(CN)_6_; 2 mM X-Gluc) was pipetted on to the detached leaf and a glass tube rolled across the leaf surface to lightly crush the tissue. Leaves were incubated overnight at 37C^0^. Prior to imaging, leaves were cleared of chlorophyll through several washes in 95% EtOH. To quantify GUS staining, multiple image processing steps were implemented using ImageJ to obtain the pixel statistics that are reported. The original RGB image was converted to HSL colorspace using the “Color Transform” plugin and the lightness channel was extracted. The image look-up table was changed to “thermal” and a manually defined circular ROI was created whose size and shape remain constant when gathering the mean and standard deviation of the pixel intensities for each of the strains. Using the same ROI, the image was cropped for each of the strains to display the exact regions sampled.

### Cassava transformation

Reporter constructs were introduced to cassava FEC cells by LBA4404 following our published methods (Chauhan *et al.*, 2015).

## Results

To shed light on the development and physiology of cassava plants from a gene expression perspective, eleven tissue/organ types from cassava cultivar TME 204 were sampled for transcriptome profiling (**Fig. 1**). Tissue type relatedness was assessed based on Jensen-Shannon (JS) distances (**Fig. 2**) and principal component analysis (PCA) (**Fig. 3**). Biological replicates clustered closely together confirming the high quality of the dataset (**Fig. S1A**, **Fig. 3A-B**). Both analyses divided the 11 tissues into three major groups: aerial (leaf, midvein, petiole, stem, lateral bud, and shoot apical meristem (SAM)), subterranean (storage root, fibrous root and root apical meristem (RAM)), and embryogenic (OES and FEC). Leaf and midvein, petiole and stem, lateral bud and SAM, and OES and FEC samples cluster together within the dendrogram (**Fig. 2B**), and occupy similar positions across the first four principal components (PCs), which collectively explain 67.3% of transcript expression level variance (**Fig. 3A-B**). These groupings are expected, representing leaf blade, vascular, shoot meristem, and callus-associated tissues. The root tissues show more complicated relationships. Figure 2 indicates storage roots as distant from fibrous roots and RAM (**Fig. 2B**). Similarly, whereas the RAM, storage root and fibrous root samples cluster closely together when projected onto PCs 1 and 2 (**Fig. 3A**), these tissues occupy more disparate positions when evaluated by PCs 3 and 4 (**Fig. 3B**). This indicates that while root samples share common gene expression patterns, tissue specific signatures differentiate storage roots from the other subterranean tissues.

**Figure 1.**
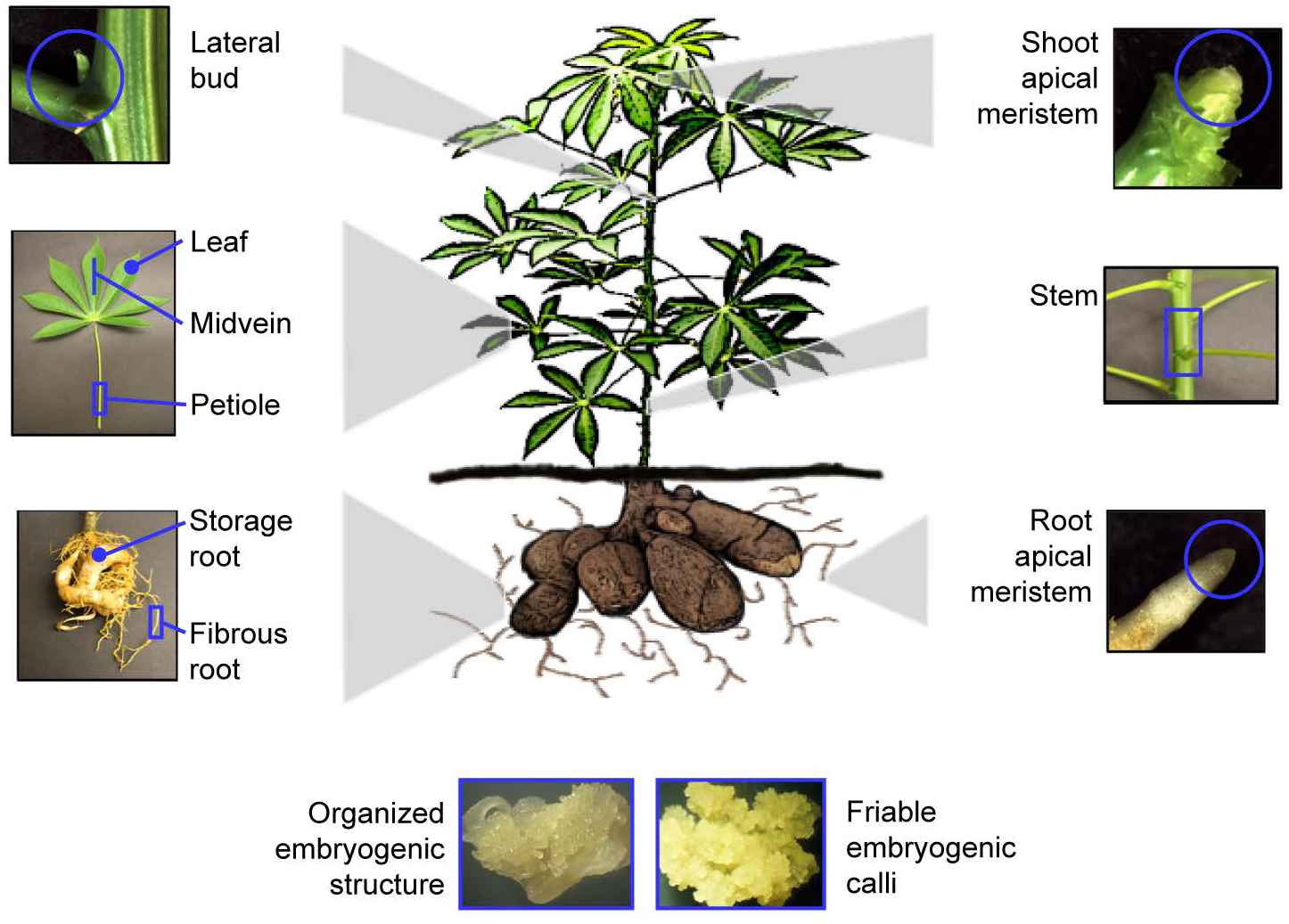
Cartoon and pictures of cassava tissues sampled for gene expression atlas. Eleven tissue types were dissected by hand and frozen in liquid nitrogen prior to processing for RNA sequencing library preparation.

**Figure 2.**
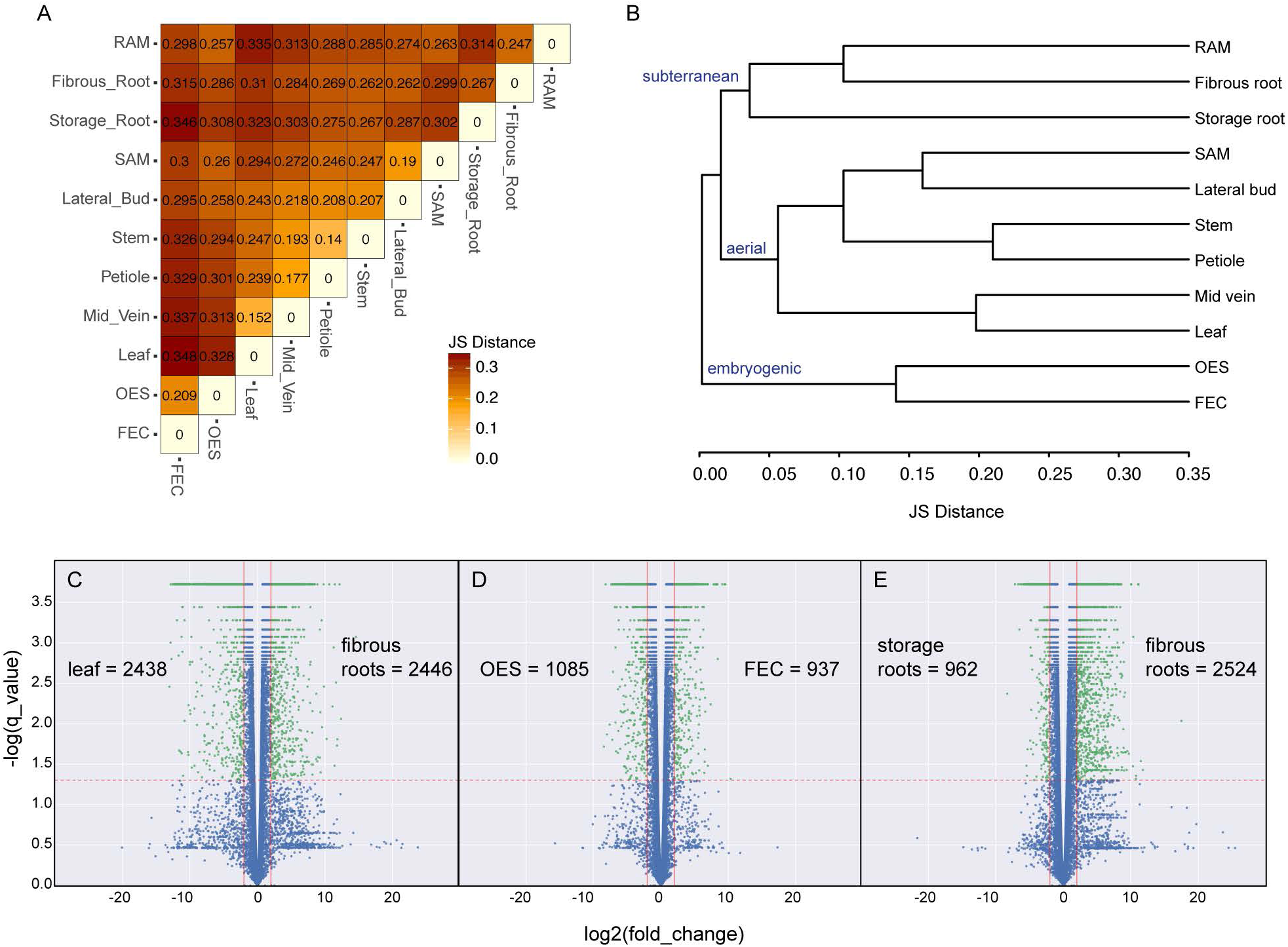
Comparison of global gene expression patterns across 11 cassava tissue types. **(A)** Heatmap showing every pairwise comparison for the 11 tissue types sampled, as produced by CummeRbund’s csDistHeat() method. Lighter colors correspond to more closely related tissue types. The numbers in each cell represent the Jensen-Shannon distance between those two tissues using the mean expression values of the biological replicates. **(B)** Output of CummeRbund’s csDendro() method. This dendrogram is created using the Jensen-Shannon distances calculated between the consensus expression values of genes for each tissue type. A low squared coefficient of variation for biological replicates was observed indicating the high quality of this dataset (Fig. S1A). **(C-E)** Volcano plots showing the differential expression of genes from leaf to fibrous roots **(C)**, OES to FEC **(D)** and storage root to fibrous root **(E)** using FDR corrected p-value as the y axis. Number of genes significantly up-regulated in each tissue type, for each comparison, are listed. Red vertical lines: +/- log2(fold_change) = 2, red horizontal line: log score = 1.3. The green points indicate significantly differentially expressed genes based on these cutoffs.

**Figure 3:**
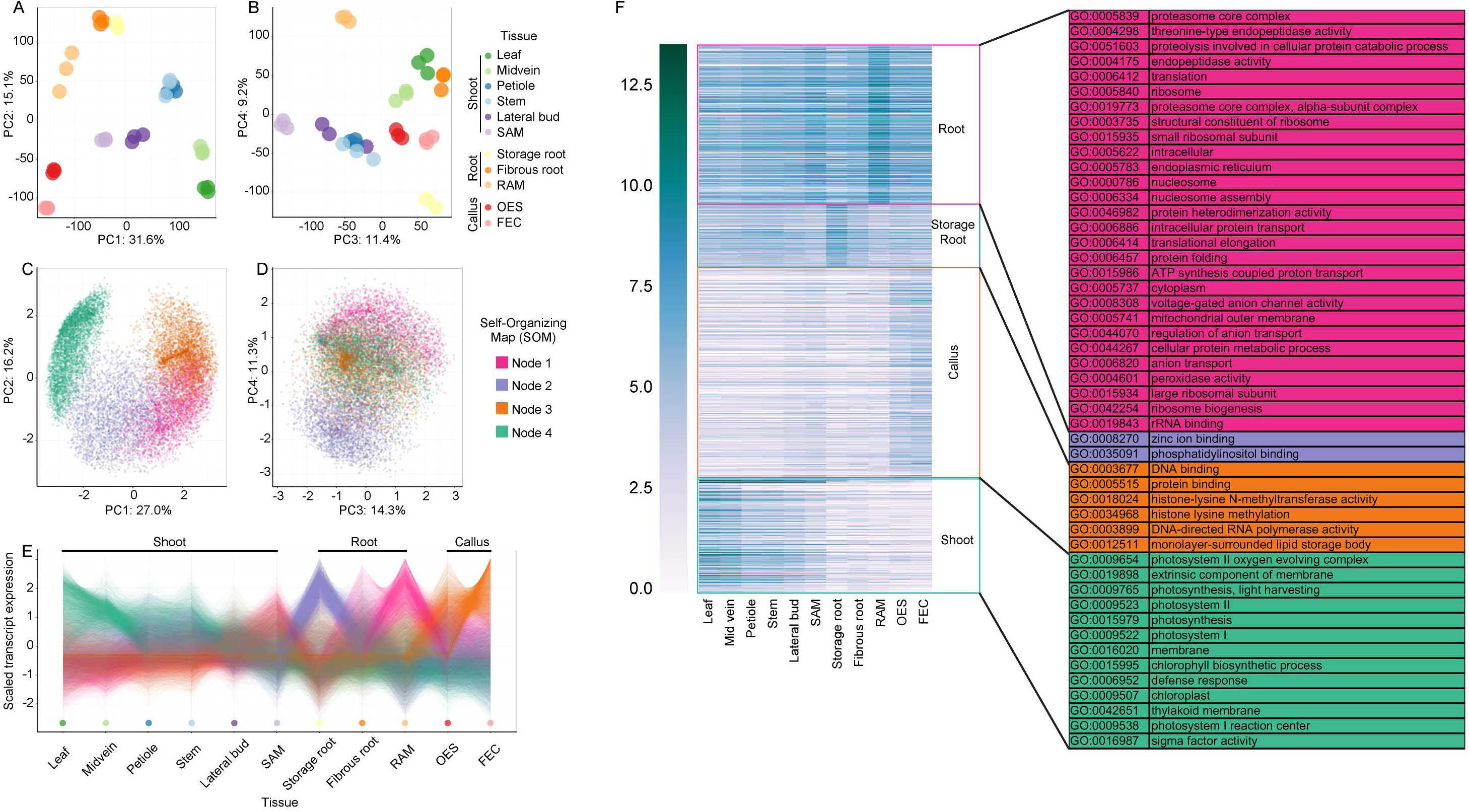
Transcript expression profiles across tissues. **(A, B)** Principal component analysis (PCA) performed on replicates of tissue samples, using transcript expression levels. **(C, D)** PCA performed on transcript profiles, across tissue samples. Colors correspond to self-organizing map (SOM) nodes used to find transcripts with similar expression profiles, which cluster together in the PCA space. **(E)** Scaled transcript expression profiles of SOM nodes across tissue types. **(F)** Heatmap of genes showing gene expression pattern corresponding to the nodes in Fig. 3C, D. Gene Ontology terms associated with these genes are listed on the right.

Two tissue comparisons within the dataset: OES vs FEC and storage vs fibrous roots, are particularly intriguing, given how little is known about the features distinguishing their physiology. The results of a PCA on the expression profiles of individual transcripts was considered. A self-organizing map (SOM) was used to identify four main clusters of transcripts with similar expression profiles across tissues, which was then projected back onto the PCA transcript space (**Fig. 3C-D**). To determine the identities of transcripts with shared expression profiles, we performed Gene Ontology (GO) enrichment analysis for each SOM node (**Fig. 3F**). In addition, we directly examined the genes most highly differentially expressed between each comparison (**Fig. 2**, **Fig. 4**).

**Figure 4.**
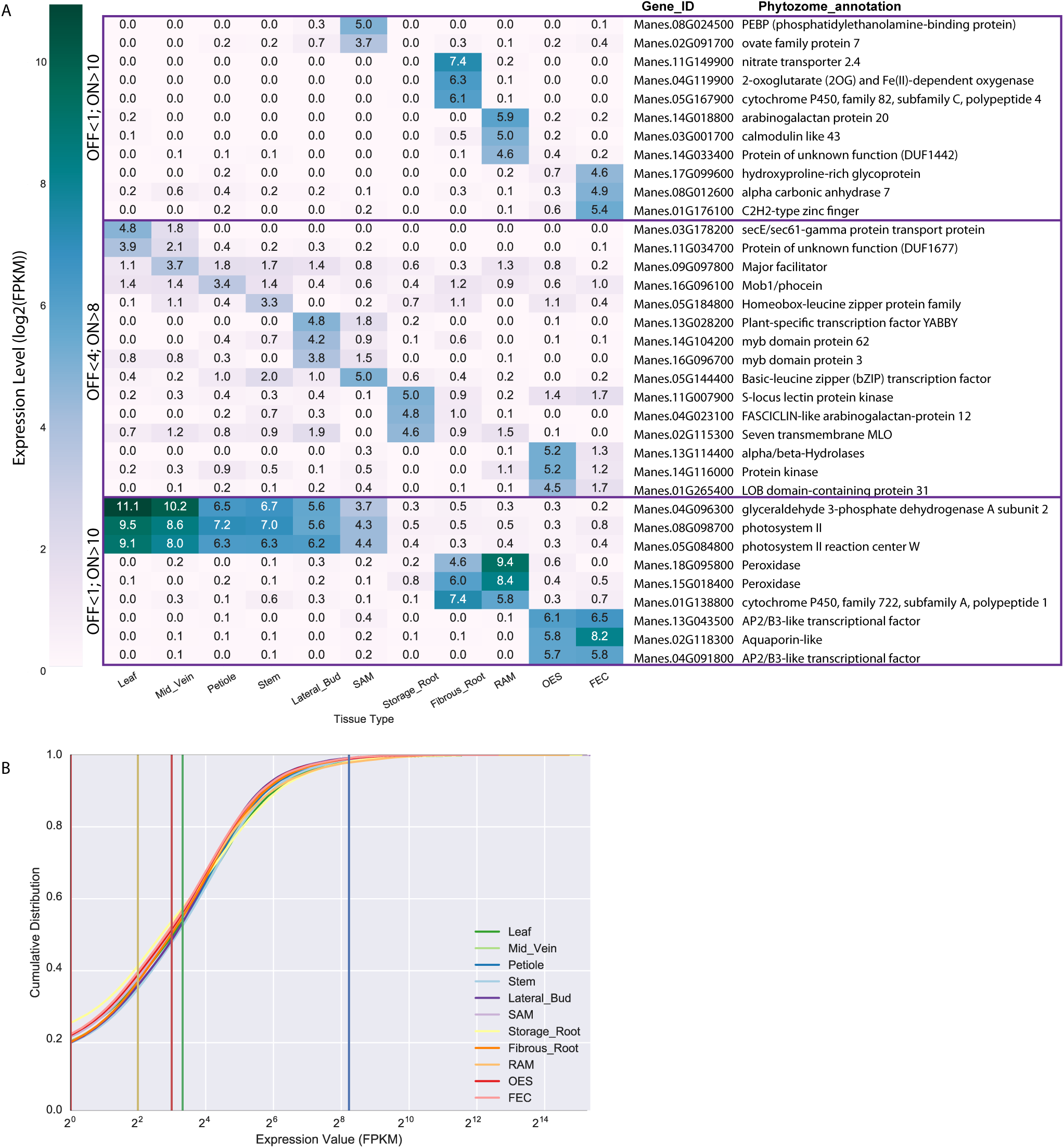
Identification of genes specifically expressed in a single or subset of tissue types. (**A**) Identification of genes with specified expression patterns. (top; middle) Heatmap of the most highly, uniquely expressed genes in each tissue. Requirements for below and above the limits of detectable expression are listed on the left (OFF and ON, respectively). (bottom) Genes expressed highly in a subset of tissues are reported. No genes specifically expressed across all subterranean tissues (storage root, fibrous root and root apical meristem (RAM)) were identified so storage root was excluded from that group. **(B)** Cumulative distribution plot of FPKM values of functionally annotated genes with expression in at least one tissue type. The vertical lines represent 3 cutoffs used in the analysis. < 1 FPKM (maroon line) = below the limit of detection; < 4 FPKM (yellow line) = below the limit of detection (relaxed); > 8 FPKM (red line) = detected expression (relaxed); >10 FPKM (green line) = detected expression; > 300 FPKM (blue line) = highly expressed genes.

First, we used the above approach to examine gene expression patterns for well characterized tissues and comparisons. Node 4 transcripts (teal) are highly expressed in the photosynthetic tissues of the leaf and mid vein (**Fig. 3E**). Similarly, comparison of leaf and fibrous roots revealed ∼4900 genes differentially expressed greater than 4-fold (|log_2_(fold_change)| > 2) between tissues (**Fig. 2C**, **SuppFile1a**). A similar number were up-regulated in each tissue and consistent with the GO term analysis presented in Figure 3F, the most highly up-regulated genes in leaf tissue pertained to photosynthesis activity while genes induced in fibrous root were related to lignin, ion binding, and transcription. We highlight that these analyses are complementary. The former takes an unbiased approach to identify variability within the dataset. The latter, directly looks at genes with maximum expression differences.

OES and FEC tissue are closely related with the latter generated from the former by a simple switch in the basal medium (Taylor *et al.*, 2012). FEC tissues are highly disorganized and ultra-juvenile in nature, consisting of proliferating, sub-millimeter sized pre-embryo units from which somatic embryos will regenerate on removal of auxin. Efficacy of FEC production from the OES is genotype dependent and can be challenging in some farmer-preferred varieties, though this recalcitrance is poorly understood (Liu *et al.*, 2011). Node 3 transcripts (burnt orange) are highly expressed in both callus tissues, but especially the FEC (**Fig. 3E**). Node 3 transcripts are associated with GO terms related to epigenomic reprogramming (DNA methylation and histone modification). Over two thousand genes were identified as differentially expressed between OES and FEC tissues (**Fig. 2D**, **SuppFile1b**). Genes up-regulated in OES tissue are associated with GO tags for heme, iron and tetrapyrrole binding and oxidoreductase activity. In contrast, genes upregulated in FEC tissue are associated with sulfur and sulfate transport (**SuppFile2**). Overall, our analyses emphasize the striking similarity between the two tissue types.

What distinguishes storage roots from other subterranean structures is ambiguous. A recent anatomical examination of these structures revealed that roots develop from the cut base of the stem cutting (basal) and from buried nodes (nodal), but that only the nodal roots will develop to form storage roots (Chaweewan & Taylor, 2015). Once initiated the storage roots develop by massive cell proliferation from the cambium to generate the central core that consists largely of xylem parenchyma in which starch is synthesized and stored. Node 1 transcripts (magenta) are highly expressed in the RAM and somewhat in the fibrous root while Node 2 transcripts (lavender) are highly expressed in the storage root suggesting that storage roots exhibit distinct gene expression patterns relative to RAM and fibrous roots. Node 1 transcripts, highly expressed in the RAM and the fibrous root, are enriched for GOs related to translation, proteolysis, and intracellular transport that might be expected for a tissue undergoing growth. Node 2 transcripts highly expressed in the storage root, are associated with zinc ion and phosphatidylinositol binding GO terms. In contrast to differentially expressed gene comparisons for leaf versus fibrous roots, and OES versus FEC, comparison of fibrous and storage roots revealed a significant shift towards gene induction in the former (**Fig. 2E**, **SuppFile1c**). Taken together, these data and analyses demonstrate that OES and FEC are highly similar tissue types and suggest that their difference may come mostly from the media on which they are cultured. In contrast, fibrous and storage roots appear as inherently distinct on a transcriptional level.

Promoters capable of driving gene expression in one or more defined tissue/organ types is essential for the successful application of biotechnology to improve crop plants. Currently, a limited set of promoters are available to achieve desired expression patterns for cassava *in planta*. For example, the root-specific patatin promoter from *Solanum tuberosum* has been used to overexpress transgenes that enhance iron and zinc levels in cassava storage roots (Gaitan-Solis *et al.*, 2015; Narayanan *et al.*, 2015). De Souza et. al. has characterized the Pt2L4 gene (Manes.09G108300) and confirmed preferential expression in cassava storage roots but also in stems (de Souza *et al.*, 2006; de Souza *et al.*, 2009). This previously published expression pattern is consistent with the current dataset. To identify cassava promoters capable of tissue specific expression, we queried the dataset for genes expressed in a single tissue type, henceforth referred to as uniquely expressed genes. To identify uniquely expressed genes, FPKM values of 1 and 10 were chosen to represent ‘below the limit of detection’ and ‘expressed’, respectively. These cutoffs were determined by investigating read mapping coverage for individual genes within our datasets. An FPKM value of less than one generally correlated with less than 1x coverage across a coding sequence. Genes expressed at greater than 10 FPKM had read mapping across the entire coding sequence. In addition, we choose an expression value of ≥300 FPKM as the cutoff for highly expressed genes which encompasses approximately the top 2% of expression values across our dataset. Below the limit of detection, expressed, and highly expressed cutoffs within the context of the entire dataset are displayed in Figure 4b. Uniquely expressed genes were identified as those expressed at greater than 10 FPKM in one tissue, and less than 1 FPKM in all other tissues (**Fig. 4a**). Applying the cutoff criteria, unique gene expression was observed for FEC, fibrous root, RAM and SAM, but not for the other seven tissues. Using less stringent cutoff FPKM values (OFF<4; ON>8), we were able to identify uniquely expressed genes for all additional tissues (**Fig. 4a**). In addition, we considered expression that would be constrained to the major groupings from the dendrogram in Figure 2. Storage root was excluded from the subterranean group because of its distinct gene expression patterns (**Fig. 2B**, **Fig. 3**).

In addition to identification of uniquely expressed genes, the data was queried to identify candidate promoters for driving strong gene expression within all surveyed tissue types (constitutive). We identified genes that showed expression values of ≥300 FPKM across our entire dataset. This analysis resulted in a list of 31 genes (**Fig. 5a**). In order to test the *in silica* analysis, promoters from five of the 31 putative constitutively expressed genes were cloned and functionally validated by fusing to the *uid*A (GUS) reporter gene. These constructs were expressed transiently in *Nicotiana benthamiana* leaves and stably transformed into cassava FEC cells. All five tested promoters were confirmed to drive GUS expression in cassava FEC cells while one promoter fusion, Manes.G035300, failed to drive expression in *N. benthamiana* for unknown reasons (**Fig. 5b**, **Fig. S2**).

**Figure 5.**
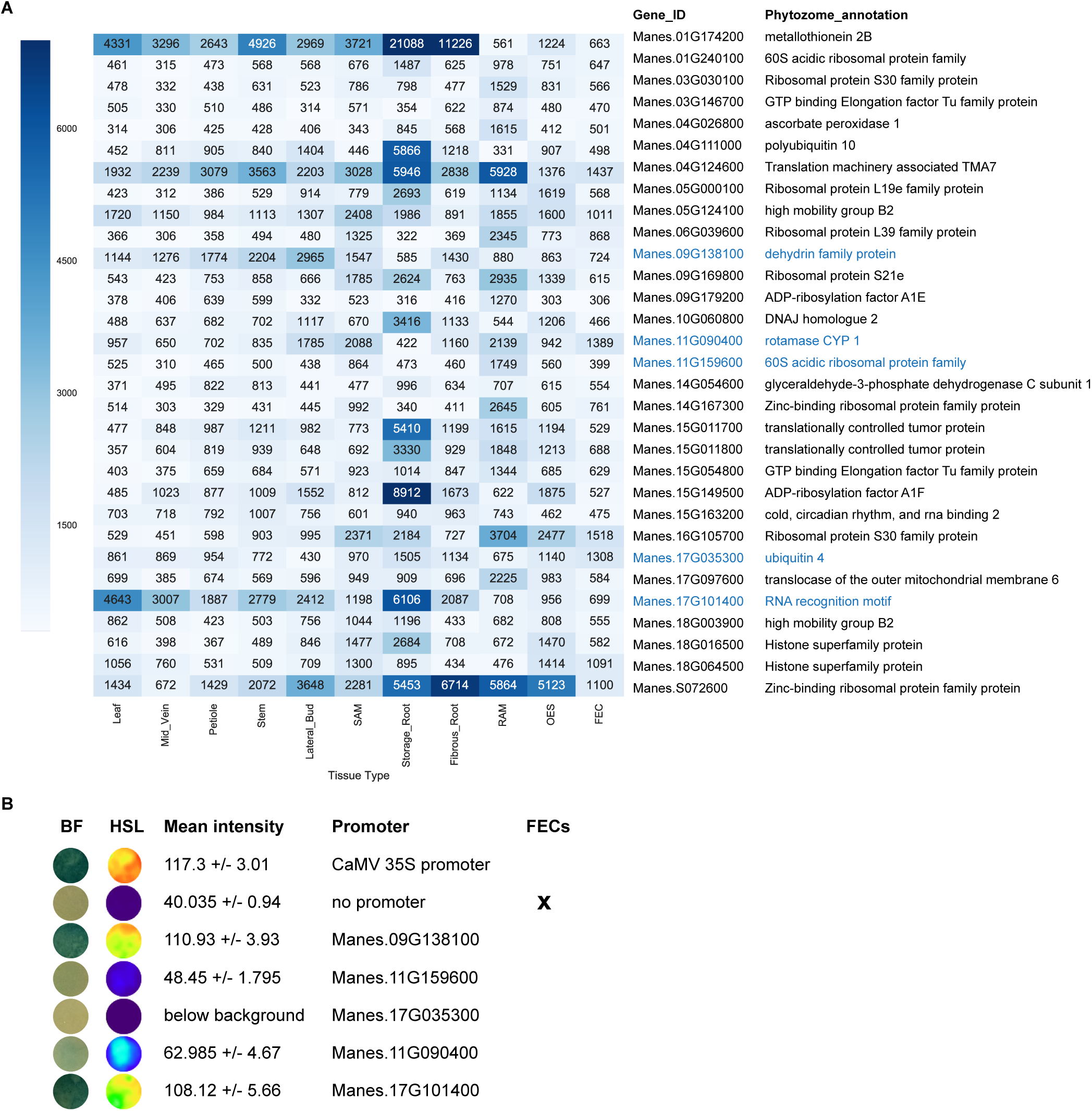
Identification of highly expressed genes in all tissue types. (**A)** Heatmap displaying expression for 31 annotated genes expressed above 300 FPKM in all tissues. Values greater than 7000 are condensed within the heatmap scale. (**B)** Ability of promoters to drive gene expression was assessed transiently in *Nicotiana benthamiana*. In addition, reporter gene constructs were stably transformed into cassava FEC cells. BF: bright field image of GUS staining intensity; HSL: images were converted to HSL color space; Mean intensity across infiltrated spot shown +/- standard deviation. Genes tested are highlighted in blue in A. Full image is shown in Figure S2.

A small collection of ‘housekeeping genes’ are routinely used for internal controls in quantitative reverse transcription polymerase chain reaction (qRT-PCR) experiments. Three cassava genes have previously been used for this purpose, *GTPb*, *PP2A*, and *UBQ10* (Moreno *et al.*, 2011). However, data from the present study show that all three genes display significant variance between tissue types (**Fig. S3**). The datasets described here were queried to identify candidate genes displaying medium level expression with low variance across the tissue types. We identified genes with expression greater than 40 FPKM in all replicates with the lowest coefficient of variation in order to normalize for magnitude of expression. Figure S3 shows the top 10 candidates from our analysis in comparison to the three genes previously used.

To facilitate future analyses, a web application has been developed wherein users can specify a desired gene expression pattern across all tissue types and receive a list of candidate genes. This application also allows users to visualize a heatmap of expression values for any gene of interest across each tissue type. The queried gene is displayed in the PCA and overlaid SOM nodes. This application can be accessed here: shiny.danforthcenter.org/cassava_atlas/

## Discussion

To assist cassava improvement efforts, various genomic, transcriptomic and epigenomic resources have previously been described (Prochnik *et al.*, 2012; Wang *et al.*, 2014; Wang *et al.*, 2015). Our study provides a unique resource: we characterize the cassava transcriptome across a wide range of tissue types. Comparison of gene expression patterns revealed a dramatic similarity between OES and FEC tissue. Storage roots were found to be significantly different from the other root tissues, and closer examination of the data suggest that the majority of this difference comes from a lack of gene expression, consistent with the role of this organ as a sink. Our study provides new insight into cassava physiology, and the data will serve as a valuable resource for cassava researchers. In addition, we identify both genes that are constitutively expressed as well as those that are highly tissue specific. The promoters of these genes may be useful for diverse biotechnological applications, including those that seek to alter cassava metabolism and improve the value of cassava as a source of food for a large fraction of the world’s population.

## Acknowledgements

This research was supported by the Bill and Melinda Gates Foundation. Sequencing was performed at the Genome Technology Access Center in the Department of Genetics at Washington University School of Medicine. The Center is partially supported by NCI Cancer Center Support Grant #P30 CA91842 to the Siteman Cancer Center and by ICTS/CTSA Grant# UL1 TR000448 from the National Center for Research Resources (NCRR), a component of the National Institutes of Health (NIH), and NIH Roadmap for Medical Research. This publication is solely the responsibility of the authors and does not necessarily represent the official view of NCRR or NIH.

### Author Contributions

M.C.W. analyzed data and co-wrote the manuscript. A.M.M. designed experiments and isolated tissues for RNAseq analysis and co-wrote the manscript. A.W.H. designed experiments and made constructs. J.B created the shiny application. R.D.C. created transgenic cassava plants. A.V. generated RNAseq libraries. N.J.T and D.F.V. supervised the study and edited the manuscript. D.H.C. performed statistical analyses and co-wrote the manuscript. R.S.B. designed experiments, supervised the study, and co-wrote the paper.

---

**Figure S1.**
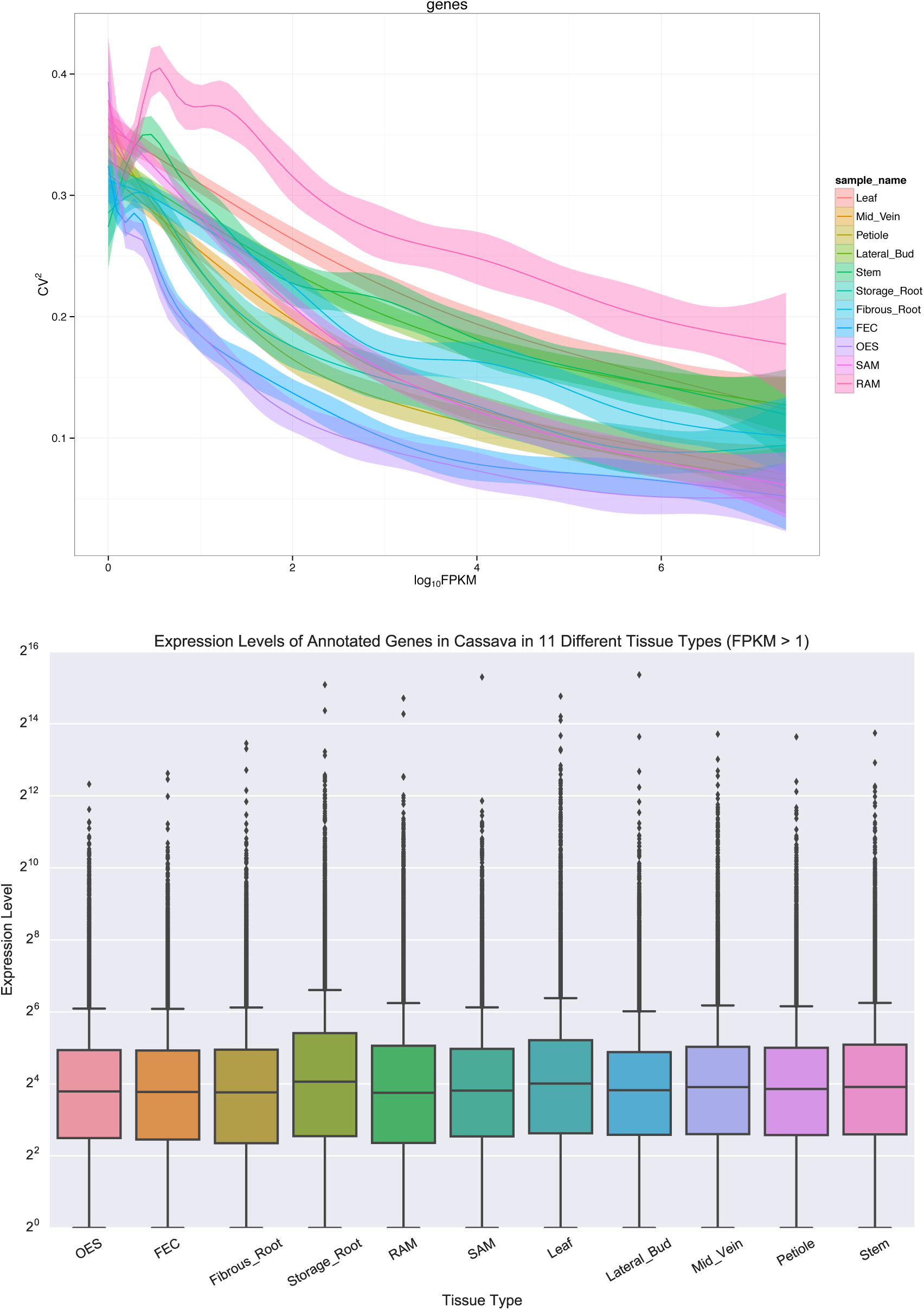
Assessing variation among biological replicates. **(A)** The squared coefficient of variation of replicates in each of the 11 tissue types is shown here to be low and reasonably uniform across all tissue types. This is indicative of replicates being closely related, limiting the possibility for error in sampling. **(B)** Distribution of all FPKM values greater than 1 in functionally annotated genes in each tissue type plotted against a log_2_ scale on the y-axis. This demonstrates the similar expression of each tissue type across annotated genes.

**Figure S2.**
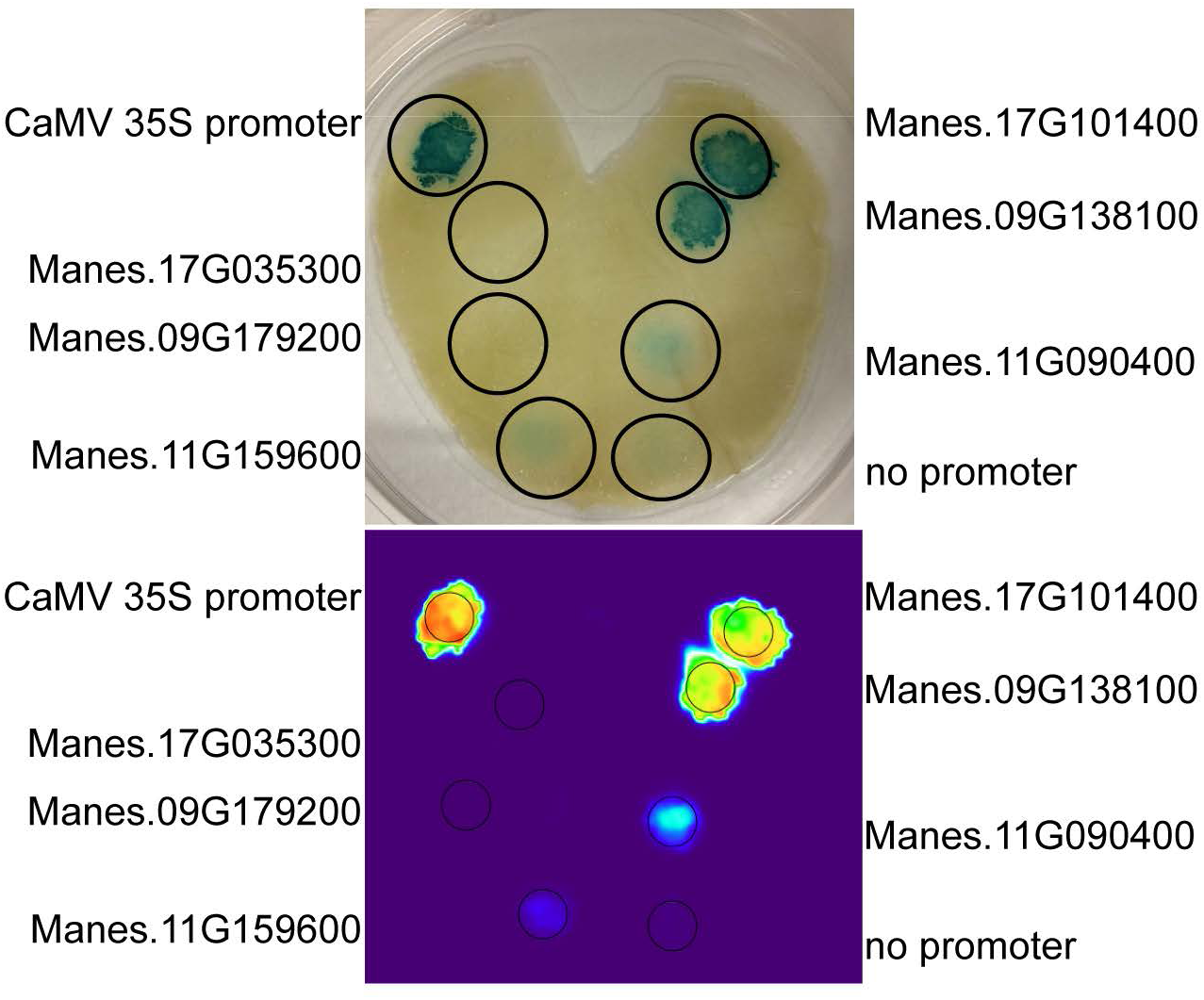
Original images that accompany Figure 5b.

**Figure S3.**
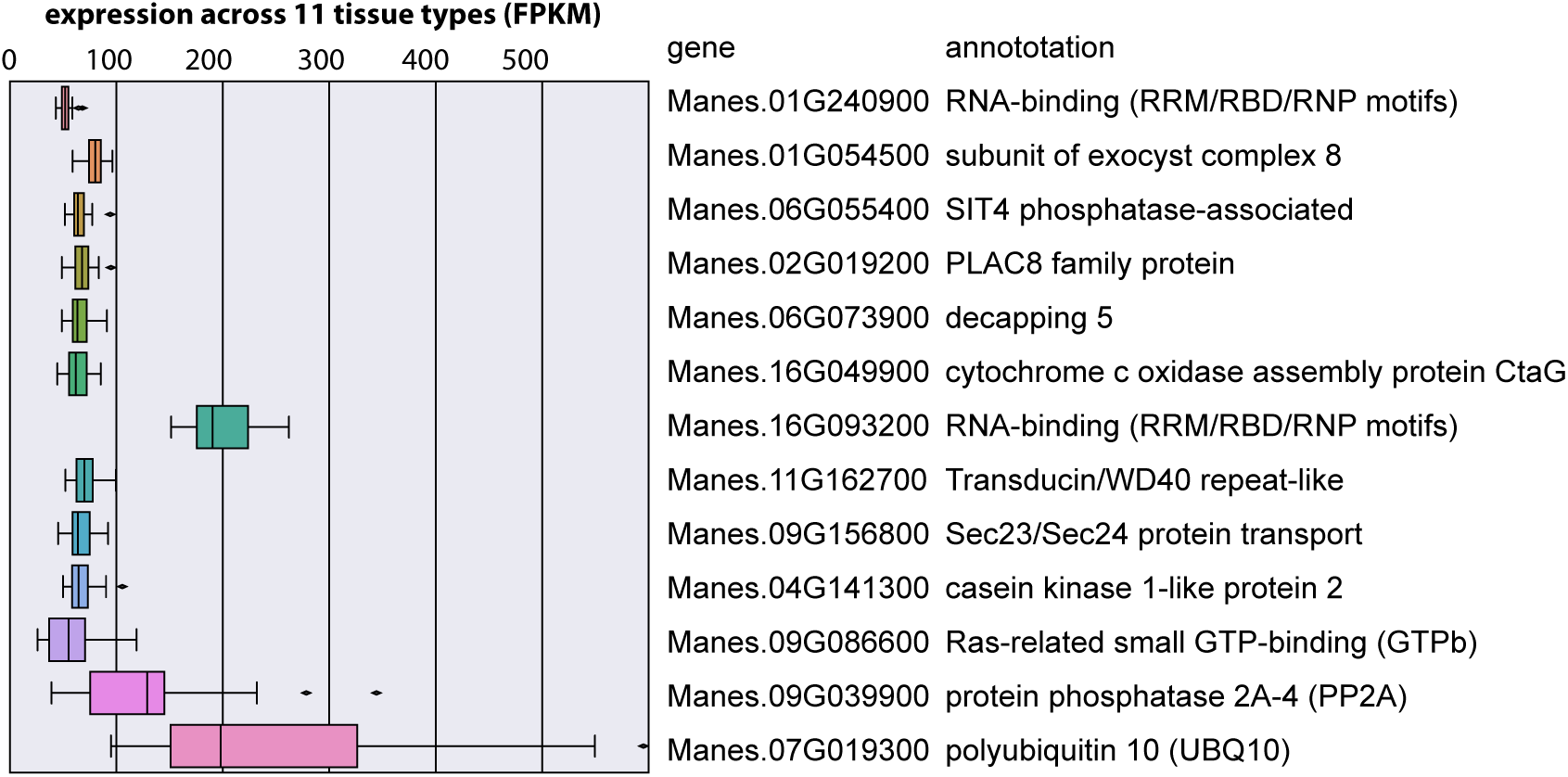
Identification of constitutively expressed genes and assessment of expression variation across tissue type. Expression profile across eleven tissue types was investigated for three housekeeping genes: GTPb, PP2A and UBQ10. The dataset was queried for genes that showed medium level expression (greater than 40 FPKM) and low variability (low coefficient of variation) across all tissues. Top ten genes are displayed.

